# Selecting Chromosomes for Polygenic Traits: Algorithms and Complexity^*^

**DOI:** 10.1101/2022.11.14.516379

**Authors:** Or Zuk

**Affiliations:** Department of Statistics and Data Science, The Hebrew University of Jerusalem, Jerusalem 91905, Israel

**Keywords:** polygenic scores, combinatorial optimization, genomic block selection, NP-hardness, semidefinite programming, synthetic genomics

## Abstract

We define and study the problem of *genomic block selection* for multiple complex traits. In this problem, one constructs a genome by selecting different genomic parts (e.g. chromosomes) from different source genomes. The constructed genome is associated with a vector of polygenic scores, obtained by summing the polygenic scores of the different genomic parts, and the goal is to minimize a given loss function of this vector. The problem is motivated by several emerging technologies: chromosome substitution lines in crop breeding, where chromosomal segments from wild relatives are combined to improve polygenic traits such as yield and stress tolerance; chromosome transfer between yeast strains for optimizing complex industrial phenotypes; and chromosomal transplantation technologies in mammalian cells. We suggest and study several natural loss functions relevant for both quantitative and threshold traits, and show that the problem is NP-complete even for a single trait and two copies, yet only weakly so, being pseudo-polynomially solvable for any fixed number of traits. We propose three algorithms with complementary roles: a Branch-and-Bound algorithm that returns the certified global optimum for any monotone loss, a fast Block-Coordinate-Descent (BCD) heuristic with random restarts that applies to any loss, and a semidefinite-programming (SDP) relaxation that provides a certified lower bound on the optimal loss for quadratic losses, and hence an optimality-gap bound when paired with the BCD solution, empirically tight in our experiments. Using the infinitesimal model for genetic architecture, we further derive, for linear losses, a closed-form approximation for the expected gain of block selection relative to random selection across multiple traits. On yeast-scale simulations BCD matches the certified Branch-and-Bound optimum on 100% of threshold-loss instances at 466× the speed, attains a certified optimality gap of at most ≈10% of the SDP lower bound for stabilizing-loss instances, and the realized gain roughly matches the analytic prediction.

## 1 Introduction

In several breeding and biotechnology contexts, it is possible or becoming feasible to combine genomic parts from different source individuals into a single genome. In plant breeding, chromosome substitution lines are routinely created by introgressing chromosomal segments from wild relatives into elite cultivars, for example barley chromosome arms into wheat [16,33], in order to improve polygenic traits such as yield, disease resistance and stress tolerance. In yeast, the completion of the Sc2.0 synthetic genome project [34] and tools for chromosome transfer between strains [31, 38] open the possibility of constructing chimeric genomes optimized for complex industrial phenotypes such as ethanol tolerance and thermotolerance, which are controlled by dozens to hundreds of quantitative trait loci (QTL) [3, 4]. In mammalian cells, high-fidelity chromosomal transplantation has been demonstrated [29, 30, 32], and programmable recombination systems now enable targeted, large-scale DNA rearrangements [12].

For highly polygenic traits where no individual gene has a large effect, selection of individual chromosomes or large-scale genomic regions from available genomes may complement local gene-editing approaches using the CRISPR-CAS9 system [21] that affect only one or a handful of genes. Chromosomal selection leverages existing genetic material without introducing new mutations, but comes with a trade-off: entire chromosomes carry both favorable and unfavorable alleles (linkage drag), whereas targeted editing can modify specific loci with precision.

Choosing which blocks to combine requires a quantitative predictor of each assembled genome’s phenotype. Polygenic scores (PS) provide this, combining the contributions of multiple genomic alleles with coefficients typically fitted from Genome-Wide-Association-Studies (GWAS). GWAS originated for human complex traits [23, 25, 40] and has since been adopted in livestock [27], crops [11], and yeast [3, 31]; accuracy continues to improve with growing sample sizes and advances in statistical methods [9, 18, 28].

Here we formulate and study the *genomic block selection* problem for multiple quantitative traits. In this problem, multiple copies are available for each chromosome (or possibly a smaller genomic part), and based on these copies’ PS we select one of them. These choices yield a genome assembled from the different chromosomal parts. We address the following two main questions:

1. How should the chromosomes be selected in order to maximize utility across multiple traits? When selecting for *T* traits, each selection **c** of chromosomes leads to an overall genomic score vector ***X***_**c**_ ∈ ℝ^*T*^ . A loss function ℒ : ℝ^*T*^ → ℝ is defined and our goal is to find the selection **c** minimizing the loss ℒ(***X***_**c**_), and compare it to the loss obtained for random selection. When *C* copies are available for each of the *M* chromosomes, the total number of possible selections *C*^*M*^ is exponential in *M* , hence the need to design efficient general algorithms for the problem.
2. What is the expected gain when using optimal chromosomal selection? How does it compare to the baseline, i.e. selecting genomes at random, as well as to the whole-genome selection procedure in which one of the *C* source genomes is selected as a whole [22, 26]?

Our main contributions are threefold. First, we formulate the genomic block selection problem mathematically and establish its weak NP-hardness even in simple cases: a single-trait stabilizing loss and a two-trait all-pass loss, each with two copies. Second, we develop three algorithms with complementary theoretical and practical roles: (i) a Branch-and-Bound (B&B) algorithm that returns the certified global optimum for monotone losses, in which the number of Pareto-non-dominated partial selections retained at each stage empirically grows much slower than exhaustive search over the regimes we tested; (ii) a simple Block-Coordinate-Descent (BCD) heuristic with random restarts, applicable to any loss, utilizing the fact that updating a single block’s selection touches only *T* coordinates of the score vector and therefore runs in *O*(*M C T*) per pass over all blocks; and (iii) a semidefinite-programming (SDP) relaxation that provides a certified lower bound on the loss for quadratic losses (empirically tight in our simulations), so it certifies the optimality gap of the BCD solution. Third, we estimate the expected gain achieved by chromosomal selection, deriving an analytic approximation for linear losses and validating the algorithms on a yeast-like simulation scenario.

## 2 The Chromosomal Selection Problem

### 2.1 Background and Selection Problems for Polygenic Scores

Consider a genome composed of *M* distinct genomic blocks, typically whole chromosomes. For haploid organisms (e.g. haploid yeast with *M* = 16), each block corresponds to a single chromosome. Diploid organisms carry two homologs per chromosome, so a block may be a chromosome type (*M*) or an individual homolog (2*M*); a diploid yeast strain, for instance, has 16 chromosome types and 32 homologs. For polyploid organisms one may further distinguish sub-genomes and homologs. For example, hexaploid wheat has 7 chromosome groups across 3 homeologous sub-genomes, so *M* can range from 7 (one block per chromosome group) through 21 (7 × 3, one block per sub-genomic chromosome) to 42 (7 × 3 × 2, one block per homolog), depending on the resolution of selection. We associate with each block a polygenic-score vector representing the genetic contribution of the block to *T* traits of interest. Suppose that we have *C* distinct source genomes, hence *C* copies are available for each block overall. Our goal is to select one copy for each block, possibly under constraints, yielding a full genome with desired properties in terms of the resulting polygenic score. Two natural selection schemes are: (i.) *whole-genome selection*, where one of the *C* source genomes is chosen, so all selected blocks belong to the same genome; (ii.) *chromosomal (block) selection*, where different blocks can be drawn from different sources, yielding a chimeric genome and a search space of size *C*^*M*^ . Concrete instances include yeast strain panels (e.g. *C* yeast strains, each with *M* = 16 chromosomes, give *C*^16^ possible chimeras) and wheat breeding (*C* donor lines, including wild relatives, provide chromosomal segments for each of the *M* chromosome groups, and one seeks the best combination of segments to optimize multiple agronomic traits). Figure 1(a) illustrates whole-genome selection, in which a single source genome is chosen, while Figure 1(b) illustrates chromosomal selection from multiple source genomes.

**Figure 1.**
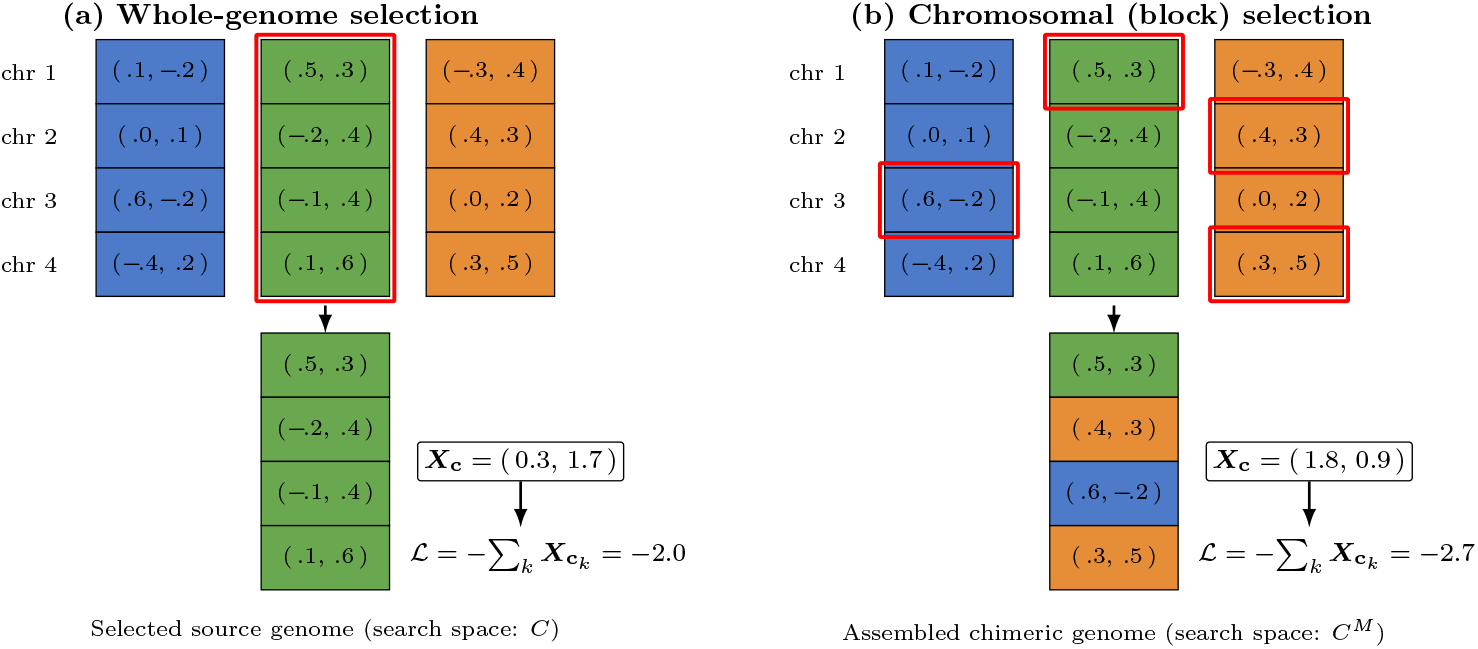
Two paradigms for selecting a genome from *C* source genomes, each providing one copy of *M* chromosomes (or genomic blocks). The example shown uses *C* = 3 source genomes (blue, green, orange), *M* = 4 chromosomes, and *T* = 2 traits, with each cell labelled by its vector of polygenic scores ***X***_*ij*•_ ∈ ℝ^2^. The two paradigms differ in which subset is selected (red boxes). **(a)** Whole-genome selection picks the single source whose aggregated polygenic-score vector ∑_*i*_ ***X***_*ij*•_ is optimal; the red box wraps around the selected genome shown as an entire column (Source 2), so the output is a monochromatic genome with ***X***_**c**_ = (0.3, 1.7). **(b)** Chromosomal (block) selection assigns, jointly across chromosomes, a source *c*_*i*_ ∈ [*C*] to each chromosome *i*, giving a chimeric genome minimizes the chosen loss (***X***_**c**_); a red box wraps around each selected block. Here the loss is the linear loss 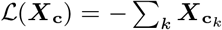 with equal weights, i.e. maximizing the total score across both traits (1.8 + 0.9 = 2.7 versus 0.3 + 1.7 = 2.0 for whole-genome selection). The output is a mosaic whose blocks are colored by their source of origin, with ***X***_**c**_ = (1.8, 0.9). Sources may represent, e.g. different yeast strains, crop donor lines, or engineered mammalian cells via chromosome transplantation.

### 2.2 Preliminaries

For a natural number *n* ∈ N denote the set {1, .., *n*} by [*n*]. For two natural numbers *m* ≤ *n* denote the set {*m, m* + 1, .., *n*} by [*m, n*]. The vector of all zeros (ones) of length *n* is denoted **0**_*n*_ (**1**_*n*_). For a polytope Δ ∈ ℝ^*n*^, define the projection operator 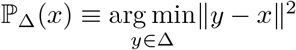.

#### Tensors

Let ***X*** ∈ ℝ_*m*×*n*×*p*_ be a 3^*rd*^-order tensor with elements *X*_*ijk*_. We denote lower-dimensional fibers of a tensor using the _•_ subscript. For example, ***X***_*ij*•_ denotes the vector (*X*_*ij*1_, .., *X*_*ijp*_) ∈ ℝ^*p*^. Similarly, ***X***_•*j*•_ is a matrix of size *m* × *p* containing all elements *X*_*ijk*_, s*i* ∈ [*m*], ∀*k* ∈ [*p*]. We also describe sub-tensors obtained by taking subsets of the indices across each dimension. For example, ***X***_[*i*][*j*]•_ is a 3^*rd*^-order tensor of size *i* × *j* × *p* obtained by taking the first *i* and *j* coordinates on the first and second dimension, respectively.

For a 3^*rd*^-order tensor ***X*** ∈ ℝ_*m*×*n*×*p*_ and a matrix **A** ∈ ℝ_*m*×*n*_, define the *2-mode* tensor-by-matrix product [1, 24], as a matrix [***X*** ×_2_ **A**] ∈ ℝ_*m*×*p*_, with elements defined by:

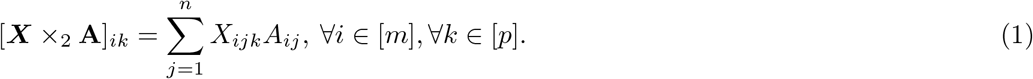

#### Gaussian distribution

For the multivariate Gaussian distribution with mean *µ* and variance Σ, denote by *ϕ*(**x**; *µ*, Σ) and Φ(**x**; *µ*, Σ) the density function and cumulative distribution function, respectively. When *µ*, Σare omitted, Φ and *ϕ* refer to the standard multivariate Gaussian distribution with mean zero and identity covariance matrix. For a measurable set *A* ⊂ ℝ^*T*^ denote by Φ(*A*; *µ*, Σ) the probability of the set under the Gaussian distribution, i.e. 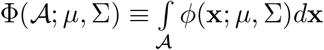.

### 2.3 Chromosomal Selection for Multiple Traits

Let ***X*** ∈ ℝ_*M*×*C*×*T*_ be a 3^*rd*^-order tensor of polygenic scores, where *X*_*ijk*_ denotes the score in chromosome *i* of copy *j* for trait *k*. Let **c** = (*c*_1_, .., *c*_*M*_) ∈ [*C*]^*M*^ be a selection vector. The associated selected polygenic vector is defined as 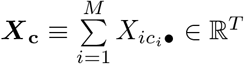 , with 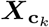 denoting its *k*-th element (***X***_**c**_)_*k*_, ∀*k* ∈ [*T*]. Our goal is to find the selected score vector ***X***_**c**_ minimizing a loss function of our choice. The *multi-trait chromosomal selection problem* is defined as follows:

#### ▶ Problem 1.

*Given a* 3^*rd*^*-order tensor of scores* ***X*** ∈ ℝ_*M*×*C*×*T*_ , *and a loss function* ℒ : ℝ^*T*→^ ℝ, *find a vector* **c**^*^ [*C*]^*M*^ *minimizing the loss:* 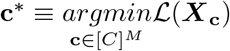.

Table 1 lists several examples of natural loss functions of interest for both quantitative and threshold traits. The computational difficulty of Problem 1 above depends on the loss function ℒ. For example, when ℒ is linear in the polygenic score vector ***X***_**c**_, we can select the optimal copy for each block independently, and the computational problem becomes trivial; this is the case when maximizing a weighted combination of quantitative traits. For non-linear losses (e.g. minimizing the overall failure probability across multiple threshold traits), selecting the best chromosome may be computationally challenging since the scores of all chromosomes must be considered jointly. In Section 3, we propose three algorithms for finding the optimal selection, applicable to general classes of loss functions.

**Table 1.**
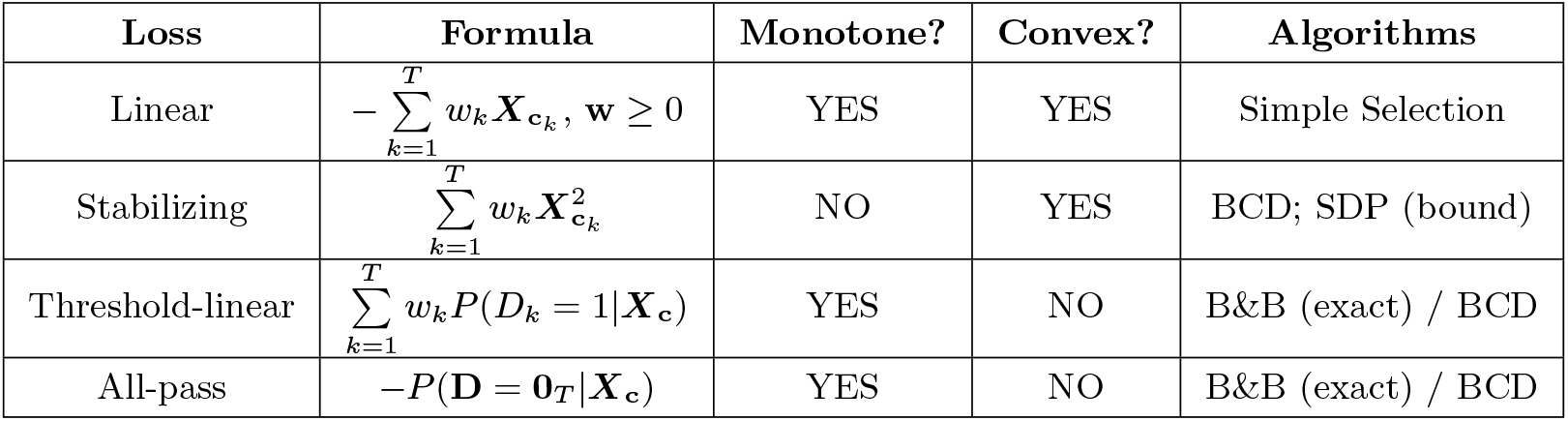
Example loss functions *L*(***X***_**c**_) and their properties. In all rows ***X***_**c**_ ∈ ℝ^*T*^ is the selected score vector and 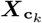 its *k*-th coordinate; **w** ∈ ℝ^*T*^ is a trait-weight vector. (i) The linear loss applies to quantitative traits (e.g. growth rate, stress tolerance, metabolite yield). (ii) The stabilizing loss [20,35,36] penalizes deviations of the resulting score from the trait optimum (set to zero w.l.o.g.); this is appropriate when a trait is already near its optimum and large deviations are harmful. (iii)–(iv) For *T* threshold traits (e.g. disease susceptibility or failure to meet a quality standard), *D*_*k*_ is a binary random variable (the status of trait *k*) whose distribution depends on the selected score ***X***_**c**_, and **D** = (*D*_1_, … , *D*_*T*_); the crossing probability *P* (*D*_*k*_ = 1 | ***X***_**c**_) follows a liability-threshold model [14] and depends on the trait’s prevalence, heritability, and the cross-trait residual covariance (full model in Appendix B). The threshold-linear loss minimizes a weighted sum of crossing probabilities; the all-pass loss takes the negative probability that no threshold is crossed.

Many natural loss functions are *monotone* in the scores vector, a property we exploit in some of our algorithms. Denote by **x** ⪯ **y** the coordinatewise product order on ℝ^*n*^ (i.e. *x*_*i*_ ≤ *y*_*i*_ for all *i*), and write **x** ≺ **y** if **x** ⪯ **y** and **x**≠ **y**. A loss ℒ : ℝ^*n*^ → ℝ is *monotonically non-increasing* if **x** ⪯ **y** implies ℒ(**x**) ≥ ℒ(**y**).

#### 2.3.1 The Gain due to Selection

##### ▶ Definition 1.

*For a tensor* ***X*** *of scores and a loss function* ℒ, *let* **c** ∼ Unif([*C*]^*M*^) *be a uniformly random selection (each c*_*i*_ *drawn i*.*i*.*d. uniformly from* [*C*]*), so that* 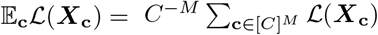. *We define the* Gain *G due to chromosomal selection and the* Gain *G*_*e*_ *due to whole-genome selection as the difference between this random-selection baseline and the optimal loss under each scheme:*

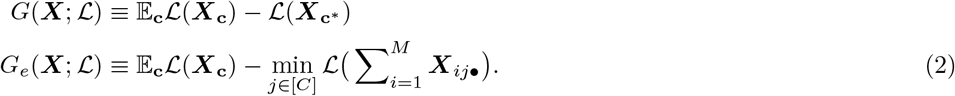

The gain *G*_*e*_(***X***; ℒ) is similar in spirit to previous definitions given in [22, 26] for human embryo selection, but generalized here for a general loss whereas [22, 26] defined the gain only for additive losses for quantitative and disease traits. By definition *G*_*e*_(***X***; ℒ) ≤ *G*(***X***; ℒ); both are functions of the score tensor ***X***. Averaging over ***X*** drawn from a statistical model yields the *expected gains* E_***X***_ *G* and E_***X***_ *G*_*e*_. In Section 4 we derive closed-form approximations of the expected gain for linear loss functions.

## 3 Algorithms and Complexity

The tractability of Problem 1 depends on the loss. For the linear loss it decomposes across blocks, each optimized independently by picking *c*_*i*_ = arg min_*j*_ −**w**^*t*^*X*_*ij*•_ in overall *O*(*M C T*) time. For general non-linear losses, we establish its computational hardness in Section 3.1. We then develop three algorithms, Branch-and-Bound (Section 3.2), Block-Coordinate-Descent (Section 3.3), and a semidefinite relaxation (Section 3.4), together with a projected-gradient relaxation [7] baseline (Section 3.4.1) that our experiments found dominated by the other methods.

### 3.1 Computational Complexity

The optimization problem 1 has an exponential search space of size *C*^*M*^ . We characterize its complexity: the next two theorems show that it is NP-hard already in simple cases: the stabilizing loss at a single trait and the all-pass loss at two traits, each with two copies.

#### ▶ Theorem 2.

*The decision version of Problem 1 with the stabilizing-selection loss, namely deciding, given a tensor* ***X*** *and a threshold τ, whether there exists* **c** ∈ [*C*]^*M*^ *with* ℒ(***X***_**c**_) ≤ *τ, is NP-complete, even for T* = 1 *and C* = 2.

**Proof**. *NP-membership:* Given **c** ∈ [*C*]^*M*^ , the score 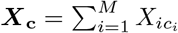 _•_ can be computed in *O*(*MT*) time and the loss evaluated in *O*(*T*) additional time, so the decision problem is in NP for any polynomial-time computable loss.

*NP-hardness:* We reduce from Partition^1^. Given positive integers *a*_1_, … , *a*_*M*_ with 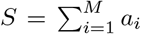 , construct an instance with *C* = 2 copies, *T* = 1 trait, and the stabilizing loss ℒ(*x*) = (*x* − *S/*2)^2^. Set the scores *X*_*i*11_ = *a*_*i*_ and *X*_*i*21_ = 0 for all *i* ∈ [*M*]. For any selection vector **c** ∈ [2]^*M*^ , the selected score is 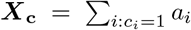, and the loss is 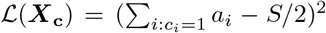. This equals zero if and only if 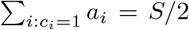, i.e. the integers {*a*_*i*_ : *c*_*i*_ = 1} form an exact partition, so a poly-time solver for our decision problem would decide Partition, hence the problem is NP-hard and therefore NP-complete.

#### ▶ Theorem 3.

*Problem 1 with the all-pass loss* ℒ(**x**) = −*P* (**D** = **0**_*T*_ |***X***_**c**_ = **x**) *is NP-hard for T* ≥ 2 *and C* = 2. *When the residual covariance is diagonal, the objective factorizes into univariate Gaussian CDFs and is efficiently evaluable*.

**Proof**. We reduce from Partition. Given positive integers *a*_1_, … , *a*_*M*_ with *S* = ∑ *a*_*i*_, set *T* = 2, *C* = 2, with scores *X*_*i*11_ = *a*_*i*_, *X*_*i*21_ = 0, *X*_*i*12_ = 0, *X*_*i*22_ = *a*_*i*_, symmetric thresholds 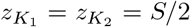, and residual covariance *σ*^2^**I**_2_ (i.e. conditionally independent trait residuals with common variance *σ*^2^), so the joint conditional probability factorises across traits. Under the liability-threshold model (eq. (13), Appendix B), each factor is the per-trait pass probability *P* (*D*_*k*_ = 0 ***X***_**c**_) = Φ (***X***_**c**_ *z*_*K*_)*/σ* , the complement of the crossing probability. For a selection with *A* = *i* : *c*_*i*_ = 1 and *u* = *a*_*i*_ we have 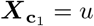 and 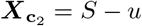, so the loss is 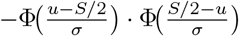. Since Φ(*z*)Φ(−*z*) = Φ(*z*)(1 − Φ(*z*)) is uniquely maximized at *z* = 0, the loss achieves its minimum −1*/*4 if and only if *u* = *S/*2, i.e. an exact partition exists.

We next show that the hardness above is only *weak*, because the problem is pseudo-polynomially solvable for any fixed number of traits.

#### ▶ Proposition 4

(Pseudo-polynomial algorithm; weak hardness). *Fix the number of traits T* , *assume integer scores X*_*ijk*_, *and let* 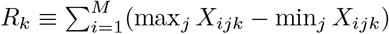 *be the range of achievable sums for trait k, with R* = max_*k*_ *R*_*k*_. *Then Problem 1 with any loss* ℒ : E^*T*^ → ℝ *evaluable in O*(*T*) *time is solved exactly in O*(*M C R*^*T*^) *time and O*(*R*^*T*^) *space. Hence for every fixed T the problem is pseudo-polynomial, so the NP-hardness of Theorems 2 and 3 is only* weak, *and bounded-precision score instances are solvable exactly in polynomial time*.

The proof, a dynamic program over the achievable score vectors, is given in Appendix A.2.

### 3.2 A Branch-and-Bound algorithm

In the Branch-and-Bound algorithm for selecting chromosomes *for monotone loss functions*, we grow a tree of all possible selected chromosomes, and at each level keep only leaves not dominated by other leaves. Finally, we evaluate the loss of all leaves at the last level. A tree of depth *b* is represented as a collection of paths from the root to the leaves Γ = {**c**^(1)^, .., **c**^(*m*)^} where each **c**^(*j*)^ ∈ [*C*]^*b*^ represents the choices of chromosomes in the first *b* levels. The partial score sum is calculated: 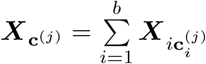 , and dominated partial score vectors are pruned.

Then, each of the remaining **c**^(*j*)^’s is expanded into *C* paths of length *b* + 1. A formal step-by-step description is shown in Algorithm 1. The computational burden of a single run is determined by the number of non-dominated partial-sum vectors retained at each stage *b*, out of *C*^*b*^ possible partial selections. In the *worst case*, the Branch-and-Bound algorithm retains all of them, hence it may run in time exponential in *M* , as shown in the Appendix, Section A.3.

While the worst-case computational complexity of Algorithm 1 is exponential in *M* , the number of vectors considered may be far lower than *C*^*M*^ in practice. Further pruning can be achieved by bounding the loss with the coordinatewise *envelopes* of the score tensor: let ***X***_*∨*_ (***X***_*∧*_) denote the vector obtained by summing, over all chromosomes *i*, the coordinatewise maximum (minimum) of *X*_*ijk*_ across *j* ∈ [*C*]. Equivalently, writing 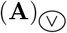 and 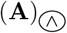 for the column-wise maximum and minimum vectors of a matrix **A** (defined in Appendix A.1 and used in line 5 of Algorithm 1), we have 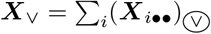 and 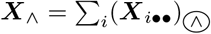 . Since ***X***_*∧*_ ⪯ ***X***_**c**_ ⪯ ***X***_*∨*_ for every feasible selection **c**, and ℒ is monotonically non-increasing, for the optimal selected score ***X***_*⋆*_ ≡ ***X***_**c***⋆*_ we obtain

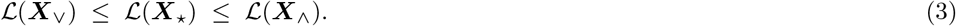

We call this an *envelope bound*: any partial-sum vector whose optimistic completion (the partial sum plus the remaining-block envelope 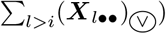 yields a loss exceeding the feasible upper bound 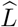 (the loss of the whole-genome optimum) cannot be optimal and is discarded. We refer to this discarding step as *envelope-pruning*.

For simulated problem instances, using the per-chromosome score model in eq. (11), the *average* number of leaves at each stage *b* was far below the *C*^*b*^ worst case over the regimes we tested (roughly *C*^*b/*2^ for *T* = 5; see Figure 2(a,b)), which makes Algorithm 1 practical for problem instances of yeast and crop scale (e.g. *C* = 2, *M* ≤ 21).

**Figure 2.**
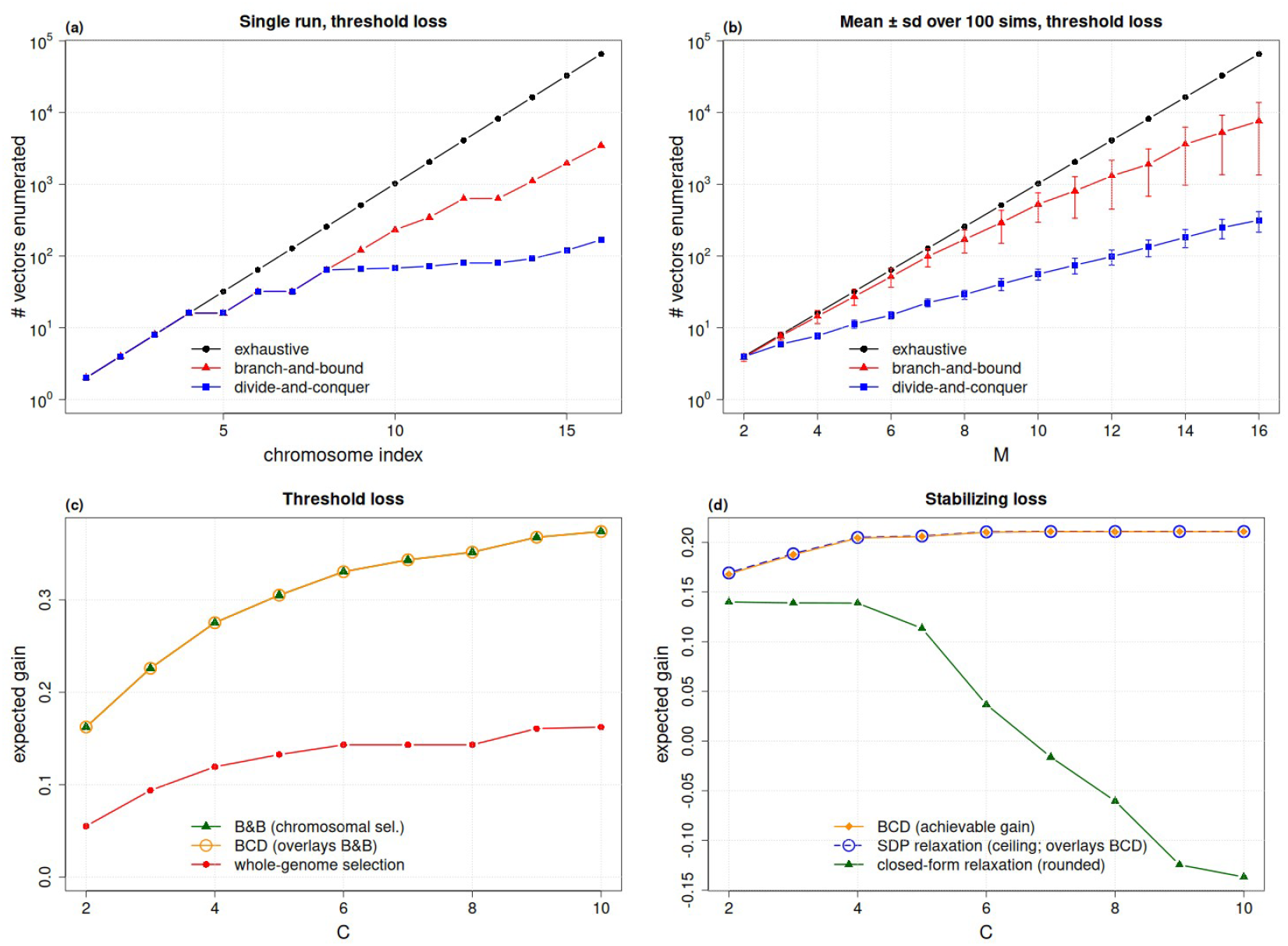
(a.) Number of Pareto-optimal partial-sum vectors retained at each stage (y-axis) vs. the chromosome index (x-axis), for a single run with *M* = 16 chromosomes (yeast), *C* = 2 source strains, and *T* = 5 traits, using the threshold loss. Black: exhaustive search (*C*^*b*^); red: Algorithm 1 (Branch-and-Bound); blue: the divide-and-conquer variant (Algorithm 4), here using *b* = 2 sub-blocks of ⌊*M*/2⌋ chromosomes each (two halves of 8 at *M* = 16). Divide-and-conquer retains *≤* 100 vectors at *M* = 16, roughly two orders of magnitude below exhaustive. (b.) Mean *±* one standard deviation of the final Pareto-set size across 100 simulations, for varying *M* ∈ [2, 16]. Both Branch-and-Bound (red) and divide-and-conquer (blue) grow much more slowly than exhaustive search over the tested regime; divide-and-conquer remains in the few-hundreds range at *M* = 16. (c.) Expected gain on the threshold loss vs. the number of source strains *C*, for yeast parameters (*M* = 16, *T* = 5, 12 simulations per *C*). Green: Branch-and-Bound (certified global optimum); orange: BCD (Algorithm 2, *R* = 10 restarts), which matched the Branch-and-Bound optimum on every simulated instance and is visually indistinguishable; red: whole-genome selection. Chromosomal selection (B&B or BCD) achieves ≈3× the expected gain of whole-genome selection. (d.) Expected gain on the (non-monotone) stabilizing loss vs. *C*, for the same parameters and 30 simulations per *C*. Orange: BCD (achievable gain); blue dashed (x markers): SDP relaxation (Algorithm 3) value, an upper bound on the achievable gain; green: closed-form Lagrangian relaxation followed by argmax rounding. BCD’s certified optimality gap relative to the SDP loss bound is 2–10% for *C* 5; on the gain scale plotted here that gap is under 1%, so the BCD (orange) and SDP (blue) curves nearly coincide. The closed-form relaxation degrades sharply and yields negative gain at *C ≥* 6.

#### Algorithm 1

hoosing Best Genome (Branch-and-Bound)

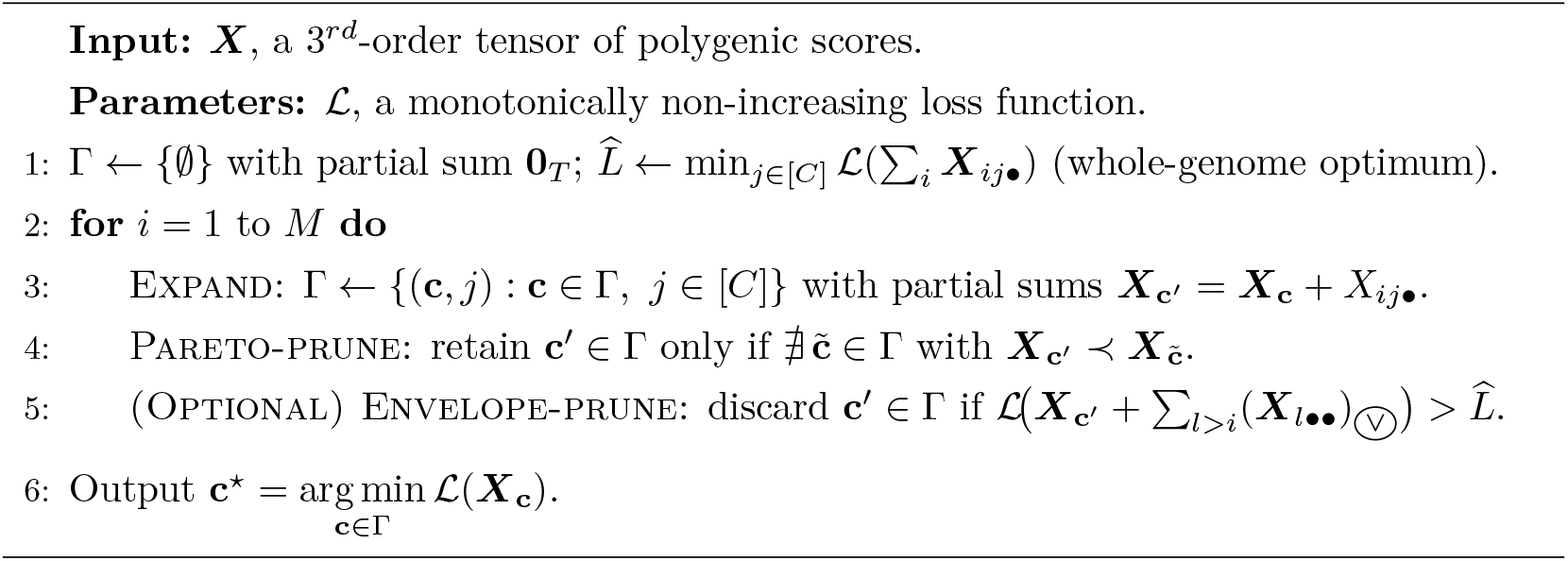

#### 3.2.1 Divide-and-conquer

We can improve the speed of our algorithm, by dividing the *M* chromosomes into groups, optimizing each of them separately, and then combining the solution in a manner where sub-vectors that cannot be extended to the optimal solution are filtered out. This procedure significantly improves performance while still being guaranteed to yield an optimal solution for monotone losses. Due to its technical details, it is described in Appendix A.4.

### 3.3 Block Coordinate Descent

Algorithm 1 (Branch-and-Bound) is inapplicable for non-monotone loss functions, and even for monotone losses its worst-case complexity is exponential in *M* . We therefore describe a simple *Block-Coordinate-Descent* (BCD) heuristic that applies to any loss and whose per-sweep complexity is linear in *M, C, T* .

The key observation is that updating a single block’s selection *c*_*i*_ while holding all other *c*_*i′*_ fixed changes only the contribution of block *i* to the aggregate score vector ***X***_**c**_ ∈ ℝ^*T*^ . Concretely, write 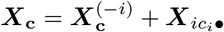 where 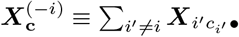 is the partial sum over all blocks except *i*. For each *j* ∈ [*C*], evaluating 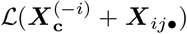 costs only *O*(*T*) once 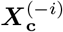 is known. A full sweep over all blocks therefore costs *O*(*M C T*) per iteration, and BCD typically converges to a 1-swap local optimum in *O*(1) sweeps. In practice we repeat the procedure from *R* random restarts and keep the best result; see Algorithm 2.

BCD has no global-optimality guarantee: it converges to a local optimum with respect to single-block swaps. However, the structural decoupling (independent blocks given the running score vector) makes it efficient enough that random restarts explore the landscape effectively in our yeast-scale simulations, where *R* = 10 restarts sufficed (Section 5).

#### Algorithm 2

Choosing Best Genome (Block Coordinate Descent)

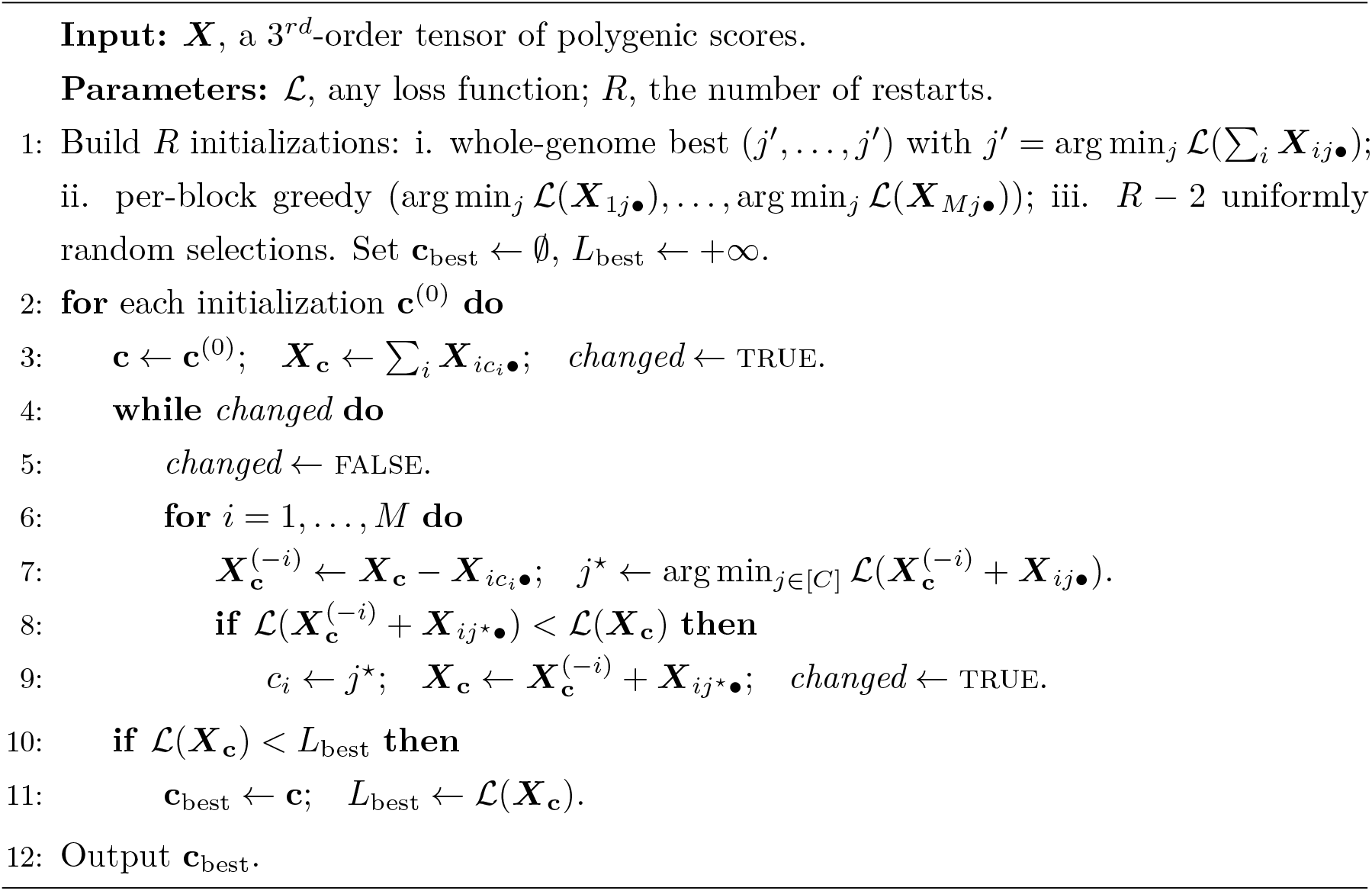

### 3.4 A Semidefinite Relaxation

For the (non-monotone) stabilizing selection loss 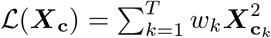 , the Branch-and-Bound algorithm of Section 3.2 does not apply and BCD has no optimality guarantee. We first review a simpler continuous relaxation that serves as a baseline, then develop a Shor-type semidefinite relaxation that provides a *certified lower bound* on the optimal loss, hence an *upper bound on the achievable gain*. Combined with a feasible heuristic such as BCD, the latter yields a certified optimality gap.

#### 3.4.1 A simpler baseline: projected-gradient relaxation

As a standard baseline we also consider a continuous-relaxation approach: encode each selection *c*_*i*_ ∈ [*C*] by a one-hot vector **C**_*i*•_ and relax **C**_*i*•_ ∈ Δ_*C*_ (the *C*-simplex); the aggregate score becomes ***X***_**c**_ = [***X*** ×_2_ **C**]^*t*^**1**_*M*_ . We minimize ℒ [***X*** ×_2_ **C**]^*t*^**1**_*M*_ over 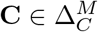 by projected gradient descent with per-row simplex projection [7] and recover an integer solution by row-wise argmax; convergence guarantees apply only when ℒ is convex [10]. For the stabilizing loss, a closed-form solution of the relaxation’s Karush–Kuhn–Tucker (KKT) optimality conditions gives a Lagrangian variant. Empirically the rounded loss was worse than the trivial baseline of picking copy 1 for every block (Section 5): the relaxed optimum sits in the simplex interior, and argmax rounding then picks essentially-arbitrary copies. This failure mode directly motivates the semidefinite relaxation below, which enforces an integer-feasible structure and yields a certified lower bound on the integer loss.

#### 3.4.2 The semidefinite relaxation

Let **q** = vec(**C**) ∈ {0, 1}^*MC*^ be the vectorized one-hot selection matrix (*C*_*ij*_ = 1 iff *c*_*i*_ = *j*; vec as defined in Appendix A.1), so *q*_(*i*−1)*C*+*j*_ = *C*_*ij*_ . The stabilizing loss is then the quadratic form ℒ(**q**) = **q**^*t*^**Aq**, where 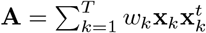 and **x**_*k*_ = vec(***X***_••*k*_). Following Shor [39], we lift the binary problem with the bordered matrix 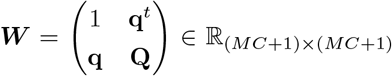 genuine selection corresponds to a *rank-*1 ***W*** ⪰ 0 with **Q** = **qq**^*t*^, so that ℒ(**q**) = tr(**AQ**); imposing this rank-1 condition, the binary constraint diag(**Q**) = **q** (valid since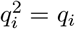for binary *q*_*i*_), and the per-block row-sum constraints reformulates Problem 1 exactly. Dropping the (non-convex) rank-1 constraint yields the SDP relaxation

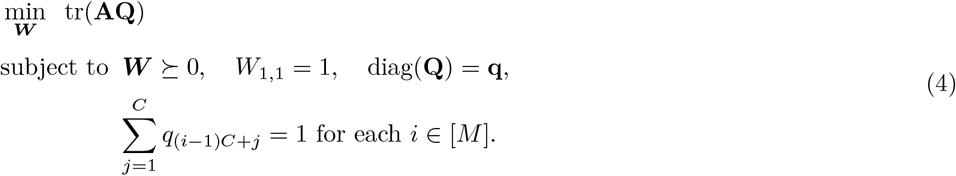

A feasible ***W*** need not be rank 1 (so the optimum may be attained at a ***W*** that is not a genuine selection); but every rank-1 integer-feasible ***W*** remains feasible, so the optimal value 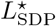 is a *lower bound* on the integer minimum loss, 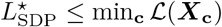 . The relaxation is solved in polynomial time using interior-point methods (we use CSDP [5]). Given any feasible integer solution **ĉ** (e.g. from BCD), the certified optimality gap is 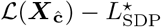 . The procedure is summarised in Algorithm 3.

##### Algorithm 3

Certified Lower Bound via Semidefinite Relaxation

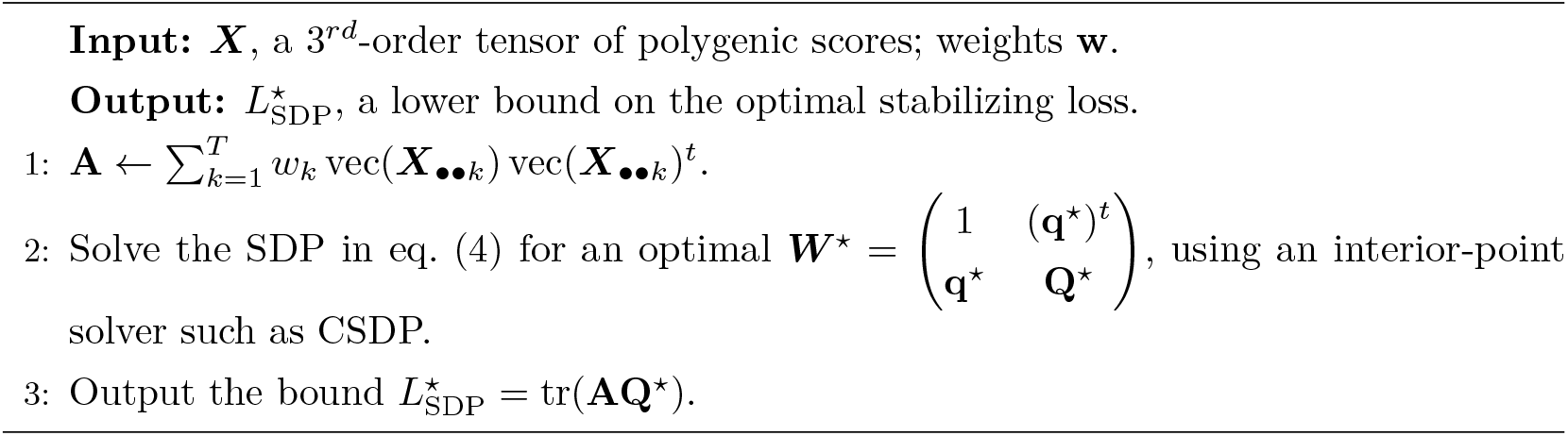

The SDP is used solely to certify the lower bound; a feasible integer selection could be recovered by block-wise argmax rounding of **q**^*⋆*^, but empirically BCD with random restarts (Algorithm 2) produced a stronger feasible solution at much lower computational cost in our experiments (Section 5), so we pair the SDP bound with the BCD solution.

## 4 Expected Gain: Analytic Prediction

To anchor the simulations that follow, we first derive a closed-form prediction for the expected gain under a population-level statistical model. Suppose the score tensor ***X*** is drawn from the matrix-Normal model of Appendix B, in which the score vector for each chromosome *i* has covariance 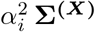 with 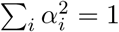, and blocks are independent. For the linear loss ℒ = −**w**^*t*^***X***_**c**_ with **w** ⪰ **0**_*T*_ and i.i.d. source genomes (**Σ**^**(*C*)**^ = **I**_*C*_), standard extreme-value arguments [22] give whole-genome and chromosomal expected gains

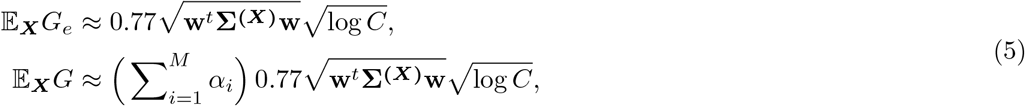

so the ratio of expected gains is ∑_*i*_*α*_*i*_. Using chromosome lengths as a proxy for 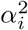 (Table 2, Appendix B), this gives a predicted ∑_*i i*_ *α*_*i*_ ≈ 3.89-fold improvement for yeast (*M* = 16) and ≈ 4.68-fold for human (*M* = 23, 22 autosomes plus the X chromosome). For non-linear losses the closed form no longer applies and we evaluate the gain numerically below.

**Table 2.**
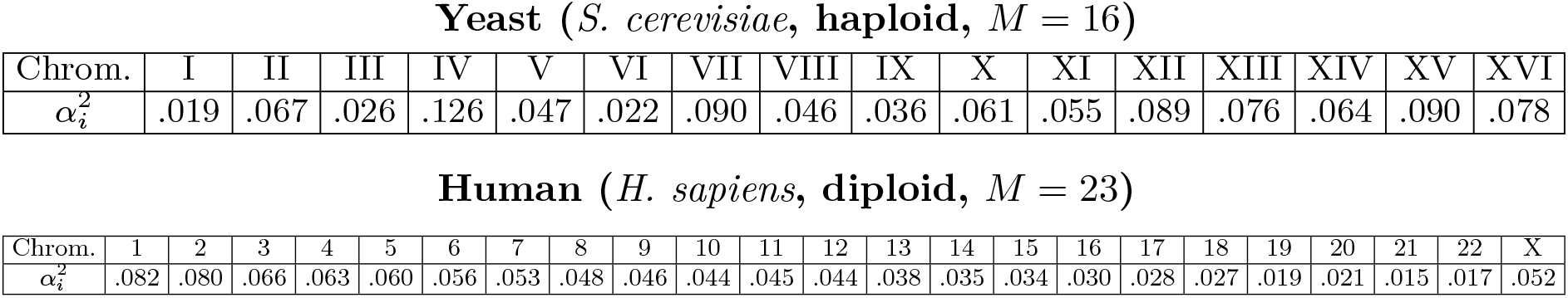
Relative chromosomal lengths (i.e. 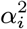 values) for the yeast (16 chromosomes) and human (22 autosomes plus the X chromosome) reference genomes. The Y chromosome contributes negligibly to autosomal polygenic scores and is omitted. For yeast, ∑_*i*_ *α* _*i*_ ≈ 3.89; for the 23-block human set, ∑ _*i*_ *α*_*i*_ ≈ 4.68.

| Chrom. | I | II | III | IV | V | VI | VII | VIII | IX | X | XI | XII | XIII | XIV | XV | XVI |
| --- | --- | --- | --- | --- | --- | --- | --- | --- | --- | --- | --- | --- | --- | --- | --- | --- |
| $\alpha_i^2$ | .019 | .067 | .026 | .126 | .047 | .022 | .090 | .046 | .036 | .061 | .055 | .089 | .076 | .064 | .090 | .078 |

■ **Table 2** Relative chromosomal lengths (i.e.
| Chrom. | 1 | 2 | 3 | 4 | 5 | 6 | 7 | 8 | 9 | 10 | 11 | 12 | 13 | 14 | 15 | 16 | 17 | 18 | 19 | 20 | 21 | 22 | X |
| --- | --- | --- | --- | --- | --- | --- | --- | --- | --- | --- | --- | --- | --- | --- | --- | --- | --- | --- | --- | --- | --- | --- | --- |
| $\alpha_i^2$ | .082 | .080 | .066 | .063 | .060 | .056 | .053 | .048 | .046 | .044 | .045 | .044 | .038 | .035 | .034 | .030 | .028 | .027 | .019 | .021 | .015 | .017 | .052 |

## 5 Simulation Results

To examine the utility of the algorithms, we have implemented them in an R package available at *github* (see Appendix C). We simulated chromosomal scores from the statistical model of Appendix B, a Matrix-Gaussian distribution [19], with parameters inspired by a yeast-like scenario: *M* = 16 chromosomes (relative lengths in Table 2), varying *C*, and *T* = 5 quantitative traits under a threshold model. Each trait has a “failure” probability of 0.1 (analogous to a disease prevalence or a probability of falling below a quality threshold), a narrow-sense heritability *h*^2^ = 50%, and polygenic scores explaining 20% of the liability variance; the residual (non-score) covariance **Σ**^**(*Y*)**^ **+ Σ**^**(*e*)**^ therefore has diagonal 0.8, with off-diagonal entries set to 0.65 (moderate cross-trait residual correlation).

We used the sum of threshold-trait probability loss function with equal weights, and the (minus) probability of passing all thresholds simultaneously. The baseline loss under random selection was, as expected, 0.1 × 5 = 0.5 for the first loss. For the all-pass criterion, the baseline pass probability *P* (**D** = **0**_*T*_) was 0.72 (so the corresponding loss is −0.72), above 0.9^5^ ≈ 0.59 that would hold under trait independence.

### Tree-pruning behavior of Branch-and-Bound (Figure 2a,b)

For a single simulated instance with *M* = 16, *C* = 2, *T* = 5, we tracked the number of Pareto-optimal partial-sum vectors retained at each stage of Algorithm 1 (Branch-and-Bound) and Algorithm 4 (divide-and-conquer). Branch-and-Bound retained ≈ 3,000 vectors at the last stage, and the divide-and-conquer variant retained only ≈ 100, roughly two orders of magnitude fewer than the exhaustive 2^16^ = 65,536 leaves. Averaging over 100 simulations for varying *M* ∈ [2, 16], the final Pareto-set size for divide-and-conquer remained in the few-hundreds range even at *M* = 16. The favorable average-case behavior can be understood through the theory of random Pareto fronts: when *C*^*b*^ i.i.d. vectors in ℝ^*T*^ are drawn, the expected number of Pareto-optimal points is *O* (log *C*^*b*^)^*T*−1^ = *O b*^*T* −1^(log *C*)^*T*−1^ [2], which for fixed *T* grows polynomially in *b*. The partial-sum vectors in Algorithm 1 are not i.i.d., but their approximate independence under the per-chromosome Matrix-Normal distribution in eq. (11) suggests a similar scaling, consistent with the empirically observed growth of ≈ *C*^*b/*2^ non-dominated vectors for *T* = 5. For *T* ≫ 5 the Pareto front grows and the tree may become prohibitively large; in that regime BCD or the SDP-bound become attractive.

### Threshold loss (Figure 2c)

For the weighted-sum threshold loss with *M* = 16, *T* = 5 and *C* ∈ [2, 10], we ran 12 independent simulations and compared the certified Branch-and-Bound optimum^2^ (Algorithm 4) against (i.) Block-Coordinate-Descent with *R* = 10 restarts (Algorithm 2) and (ii.) whole-genome selection (min_*j*∈[*C*]_ ℒ(∑_*i*_ ***X***_*ij*•_)). BCD matched the Branch-and-Bound optimum on all 108 simulated instances (12 replicates per *C, C* ∈ [2, 10]); mean BCD running time was 0.01 s vs. 4.8 s for Branch-and-Bound at *C* = 10 (all timings in this section use the divide-and-conquer variant, Algorithm 4); summed over all *C* ∈ [2, 10], BCD’s total running time was 466× shorter (the per-*C* speed-up grows with *C* as Branch-and-Bound slows). To confirm these conclusions are not specific to *M* = 16, *T* = 5, we repeated this comparison across *M* ∈ {8, 16, 23} and *T* ∈ {2, 3, 5} (*C* = 5, 10 replicates each; 90 instances); BCD with *R* = 10 restarts again matched the certified Branch-and-Bound optimum on every instance. Branch-and-Bound time grew with problem size, reaching ≈1.7 s at *M* = 23 and over an hour per instance at *T* = 6 (as the Pareto front expands), while BCD stayed ≈0.01 s throughout. Chromosomal selection achieved roughly 3× the expected gain of whole-genome selection at every value of *C* tested. This empirical ratio is consistent with the analytic prediction of _*i*_ *α*_*i*_ ≈ 3.89 for yeast derived in eq. (5) for the linear loss: although the threshold loss is non-linear, over the simulated range it behaves approximately linearly in the relevant score directions, giving a similar block-additive geometry, so the ratio of expected gains tracks the linear prediction closely. The full agreement between BCD and B&B suggests that instances drawn from the matrix-Normal model in eq. (11) induce a benign optimization landscape for BCD, where *R* = 10 restarts suffice with high probability. On adversarial inputs (e.g. Partition-style constructions, cf. Theorem 2) BCD’s 1-swap neighbourhood is no longer sufficient and the worst-case gap can be large. The projected-gradient baseline (Section 3.4.1) was again worse than the trivial copy-1 selection and is not displayed in Figure 2c.

### Stabilizing loss (Figure 2d)

For the (non-monotone) stabilizing loss with the same yeast parameters and 30 simulations per *C*, we computed (i) the BCD solution (Algorithm 2) and (ii) the SDP-guaranteed lower bound (Algorithm 3). Across all 270 simulated instances (30 replicates per *C, C* ∈ [2, 10]) the SDP bound was below BCD’s feasible loss. Averaging across replicates, BCD’s certified optimality gap, the relative excess of its feasible loss over the SDP lower bound 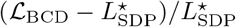 , was 2.5%, 4.1%, 10.1% and 8.6% at *C* = 2, 3, 4, 5, respectively; at *C* ≥ 6 the SDP bound and the BCD loss both approach zero and the absolute gap shrinks below 10^−3^. Note that Figure 2(d) plots the resulting expected *gain*, on whose scale this gap is under 1%, so the BCD and SDP-ceiling curves are visually indistinguishable. In contrast, the closed-form Lagrangian relaxation (dropping the nonneg constraint, solving the row-sum-constrained quadratic in closed form, then rounding by row-wise argmax; plotted in green) degrades sharply with *C* and yields negative gain (worse than the trivial baseline) at *C* ≥ 6, illustrating the failure mode of simplex-relaxation+rounding for this loss.

### Practical recommendation

Based on these results, we recommend BCD with random restarts as the default algorithm in practice. Branch-and-Bound (Section 3.2) should be used when a certified global optimum is required for a monotone loss, and is tractable at the yeast/crop scale (*M* ≲ 21, *C* ≲ 10); at larger scale (e.g. when blocks are taken to be finer-grained chromosomal segments, *M* ≫ 21, or when the Pareto front blows up at large *T*), the certified branch is infeasible and BCD becomes the only practical option. The SDP lower bound (Section 3.4) provides an optimality-gap certificate for quadratic losses; its complexity grows as (*MC*)^*O*(1)^ via interior-point methods, so it remains usable beyond the regime where B&B applies, though it too eventually scales out and one must fall back on BCD alone.

## 6 Discussion

We have defined and formulated the genomic block selection problem and provided three algorithms with complementary roles. Our Branch-and-Bound algorithm (Algorithm 1) returns the certified global optimum for any monotone loss; despite worst-case exponential complexity, the divide-and-conquer variant (Algorithm 4) kept the retained Pareto set in the few-hundreds range at *M* = 16, while the plain version retained ≈ 3,000 vectors, both well within reach for yeast and crop scenarios (*M* ≤ 21). The Block-Coordinate-Descent heuristic (Algorithm 2) applies to any loss, runs in milliseconds on yeast-scale instances, and was empirically optimal on every threshold-loss instance we tested (all 108 simulated instances). For the non-monotone stabilizing loss the SDP relaxation (Algorithm 3) provides a certified lower bound that, combined with the BCD solution, yielded a certified optimality gap of ≲ 10% for *C* ≤ 5 in our simulations. The standard projected-gradient relaxation (Section 3.4.1), included as a baseline, was strictly dominated by BCD on both monotone and non-monotone losses in our setup. Establishing problem-instance-dependent approximation guarantees for BCD and tightening the SDP at larger *C* are interesting directions for future research.

The problem is motivated by several current and emerging technologies. In crop breeding, chromosome substitution lines from wild relatives are routinely used to improve polygenic traits in wheat and other polyploid crops [16, 33], yet the optimal combination of donor segments for multiple traits is typically chosen by breeder intuition rather than formal optimization; our algorithms apply directly once polygenic scores are available. In yeast, the Sc2.0 synthetic genome project [34] and panels of sequenced strains [31] make chimeric genomes for industrial phenotypes feasible (across dozens of yeast growth traits, including high-temperature growth, mapped QTL explain a median of ≈ 73% of the additive heritability [3]). In mammalian cells, high-fidelity chromosome transplantation has been demonstrated [32], suggesting future relevance to animal models and potentially human applications, subject to technological and ethical constraints.

Our framework assumes additive polygenic scores across chromosomes, a consequence of the infinitesimal model; cross-chromosome epistasis, particularly between divergent source genomes, could reduce the realized gain, so eq. (5) should be read as an optimistic benchmark under the additive model rather than a worst-case bound. Our empirical evaluation, moreover, uses scores simulated from the matrix-Normal model (calibrated to real yeast chromosome lengths and heritabilities); validating the algorithms on empirical per-chromosome polygenic scores (e.g. from sequenced yeast strain panels or crop populations) is an important next step. Extending the framework to higher-order chromosomal interactions, finer-grained selection units (chromosomal segments rather than whole chromosomes), and nonlinear polygenic scores are all important directions for future work.

## Appendix

### A Algorithms and Optimization Details

#### A.1 Notations

We use a single *block-by-block* vectorization throughout the paper. For a matrix *X* ∈ ℝ_*m*×*n*_, vec(*X*) ∈ ℝ^*mn*^ is defined by [vec(*X*)]_(*i*−1)*n*+*j*_ = *X*_*ij*_ (stacking the rows 1, … , *m* in order). Similarly, for a 3^*rd*^-order tensor *X* ∈ ℝ_*m*×*n*×*p*_, mat(*X*) ∈ ℝ_*mn*×*p*_ has row (*i* − 1)*n* + *j* equal to the fiber *X*_*ij*•_.

There are 2^*T*^ possible binary vectors of length *T* , with each such vector **d** ∈ {0, 1}^*T*^ , corresponding to an orthant *O*_**d**_ ⊂ ℝ^*T*^ defined as 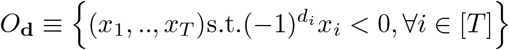 . These 2^*T*^ orthants form a disjoint union of ℝ^*T*^ (ignoring equalities with the axes).

In similar to eq. (1), for a vector **V** ∈ ℝ^*p*^, define the *3-mode* tensor-by-vector product as the matrix [***X*** ×_3_ **V**] ∈ ℝ_*m*×*n*_, with elements defined by:

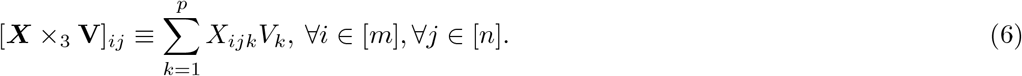

##### Element-wise notations

For two vectors or matrices *A, B* of the same size, their Hadamard product *A* ⊙ *B* is the entrywise product (e.g. [*A* ⊙ *B*]_*ij*_ = *a*_*ij*_ *b*_*ij*_ for matrices). For a matrix *A*, the column-wise maximum and minimum vectors are denoted 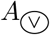 , 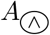 , with 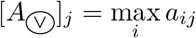 and 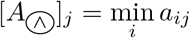 (the max/min over rows, for each column).

#### A.2 Proof of the pseudo-polynomial algorithm

**Proof of Proposition 4**. Shift each trait so that min_*j*_ *X*_*ijk*_ = 0 for every *i, k*; this translates every achievable score vector, and the loss argument, by a fixed vector, leaving the minimizer unchanged. Every partial score vector then lies in the grid 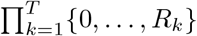. Let *S*_*i*_ be the set of partial-sum vectors attainable using blocks 1, … , *i*: *S*_0_ = {**0**_*T*_ } and *S*_*i*_ = {***v*** + *X*_*ij*•_ : ***v*** *S*_*i*−1_, *j* ∈ [*C*]} . Each *S*_*i*_ is a subset of the grid, so |*S*_*i*_|*∏*_*k*_ (*R*_*k*_+1), and computing *S*_1_, … , *S*_*M*_ costs *O*(*M C∏*_*k*_ *R*_*k*_); returning min{ℒ (***v***) : ***v*** *S*_*M*_ gives the exact optimum. As *∏*_*k*_ *R*_*k*_ is polynomial in the largest score magnitude for fixed *T* , this is a pseudo-polynomial algorithm. ▪

#### A.3 Worst-case behavior of Branch-and-Bound

##### ▶ Proposition 5.

*In the worst case, the number of non-dominated vectors at stage b of Algorithm 1 is C*^*b*^.

**Proof**. We construct an instance with *T* = 2 traits as follows. Draw 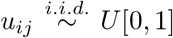for *i* ∈ [*M*], *j* ∈ [*C*], and set *X*_*ij*•_ ≡ (*u*_*ij*_ , −*u*_*ij*_). For any selection **c** ∈ [*C*]^*b*^, the partial sum is 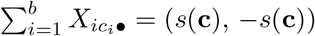, where 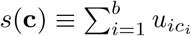 . With probability 1 over the draws, the *C*^*b*^ values {*s*(**c**) : **c** ∈ [*C*]^*b*^} are all distinct. For two selections **c**≠ **c**^*′*^ with *s*(**c**) *< s*(**c**^*′*^) the partial sums (*s*(**c**), −*s*(**c**)) and (*s*(**c**^*′*^), −*s*(**c**^*′*^)) are coordinatewise incomparable: the first has a smaller first coordinate but a larger second coordinate. Hence all *C*^*b*^ partial sums are pairwise Pareto-incomparable and Algorithm 1 retains all of them at stage *b*.

#### A.4 The divide-and-conquer algorithm

Partition the *M* chromosomes into *b* disjoint sub-blocks *B*_1_, … , *B*_*b*_ with ⊍_*i*_*B*_*i*_ = [*M*] and run Algorithm 1 on each sub-tensor 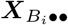 to obtain the Pareto-optimal set Γ^(*i*)^ and sub-block optimum 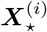. Writing 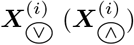 for the coordinatewise max (min) over Γ^(*i*)^, the global optimum satisfies 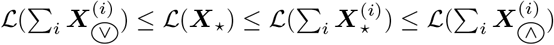 for any non-increasing ℒ. The first inequality gives a lower-bound certificate; the second gives the monotonically best loss found so far, 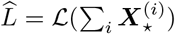, used for envelope pruning (line 7 of Algorithm 4). Algorithm 4 combines per-sub-block enumeration with Pareto- and envelope-pruning during the merge phase.

##### Algorithm 4

Choosing Best Genome (Branch-and-Bound Divide-and-Conquer)

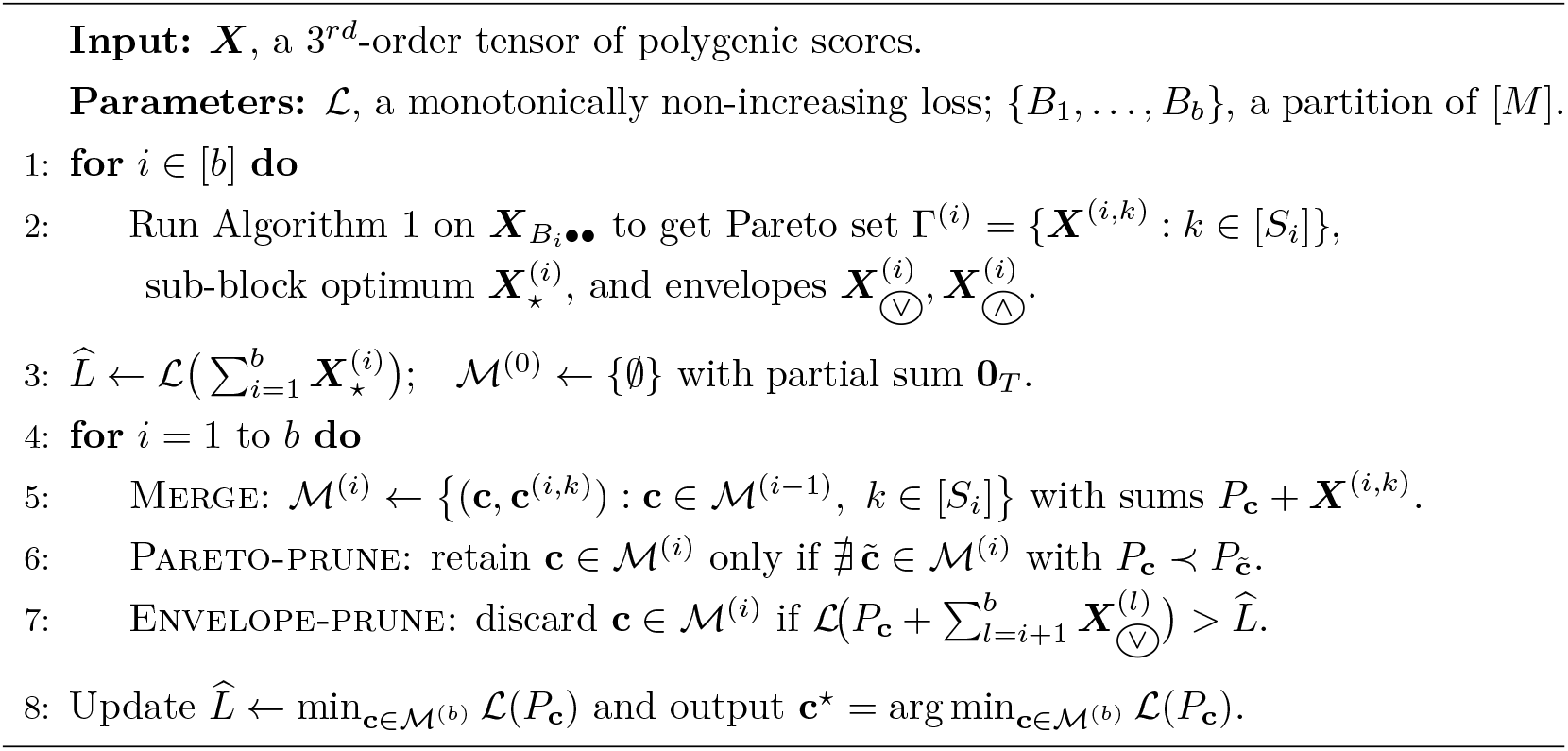

▶ Remark 6 (Correctness of Algorithm 4). For monotonically non-increasing ℒ and any feasible extension **c**^*′*^ of a partial selection **c** on the first *i* sub-blocks, the partial sum satisfies 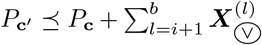 coordinatewise, hence 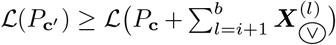. If this lower bound exceeds the best loss found so far, 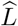, no extension of **c** can improve on it, and the envelope-pruning step in Algorithm 4 is sound. Together with the Pareto-pruning step it follows that the final selection **c**^*⋆*^ is globally optimal.

#### A.2 Computing the Gradient

We show as an example the gradient computation for the stabilizing-selection loss and the threshold-linear (disease) loss. We also show that the simplex relaxation of Problem 1 (Section 3.4.1) is convex for the first case, and not convex for the second case.

1. For the stabilizing selection loss 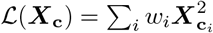, the relaxed problem is convex in **C** (the Hessian is a sum of rank-1 PSD outer products 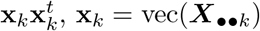, weighted by *w*_*k*_ *>* 0), and its gradient and closed-form KKT solution are direct consequences of the quadratic form; the SDP relaxation of Section 3.4 provides a strictly tighter bound and is the route we follow in this paper.
2. For the threshold-linear loss ℒ= ∑_*i*_ *w*_*i*_ *P* (*D*_*i*_ = 1 ***X***_*c*_) with 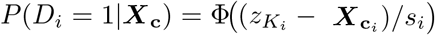 and residual standard deviations 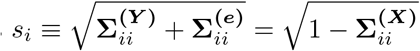 (using the unit marginal-variance normalization), the gradient is

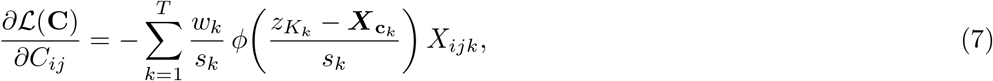

with matrix form ∇ℒ(**C**) = − ***X*** ×_3_ [**w** ⊙ ***s***^−1^ ⊙ *ϕ* (**(*z***_*K*_ − ***X***_**c**_) ⊙ ***s***^−1^)] , where 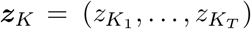 and 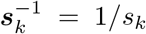 . Differentiating once more gives *∂*^2^ℒ*/∂C*_*ij*_ *∂C*_*mc*_ = − ∑_*k*_ β_*k*_ *X* _*ijk*_ *X* _*mck*_ 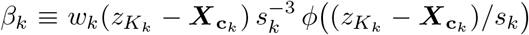 whose sign flips as 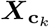 crosses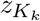 ; the Hessian is therefore not positive semidefinite uniformly over **C** and the loss is non-convex.

### B A Statistical Model from Quantitative Genetics

To put the abstract problem of the previous section in context, we describe a quantitative-genetics model for the joint distribution of the scores and non-score components determining a phenotype across *C* genomes and *T* complex traits. The model is general, applying to any organism for which polygenic scores can be computed, and it extends models used in [22, 26] for multiple traits, with two main differences: First, we model explicitly the joint distribution of the individual chromosomes’ scores, whereas in [22, 26] a model was given for the entire genomic score. Second, [22, 26] considered embryos derived from the same two parents, yielding a specific genetic relationship matrix Σ^(*C*)^ representing the Identity-by-Descent sharing of siblings, while in our case the genetic relationship matrix may be more general depending on the selection scenario (e.g. unrelated yeast strains or crop varieties, for which **Σ**^**(*C*)**^ ≈ ***I***_***C***_).

We assume that the genetic architecture of the traits is infinitesimal, namely that there are numerous causal variants, uniformly distributed along the genome. Denote the matrix of quantitative trait values as **Z** ∈ ℝ_*C*×*T*_ , where *Z*_*ij*_ denotes the value of trait *j* for the *i*-th copy. We can decompose **Z** as follows:

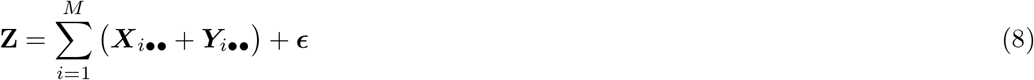

where the error term ***Y*** represents a tensor of genetic components not accounted for by the scores ***X***, and the error term ***ϵ*** represents a matrix of environmental components, both having zero mean.

We assume that all the traits have mean zero and variance 1, and further that the individual chromosome scores also have zero mean.

We further assume that the distribution of the polygenic scores ***X*** is approximately Normal in each genome (due to the polygenic nature of most complex traits [40]), and that the joint distribution of the polygenic scores over *C* genomes is multivariate Gaussian.

Consider *T* traits normalized to have zero mean and unit variance. 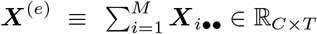 be the entire-genome score matrix for the *C* copies, obtained by summing the per-chromosome score matrices (superscript *e* for aggregation over all blocks, as in *G*_*e*_; not the environmental **Σ**^**(*e*)**^), and similarly 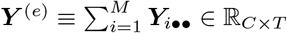 . The vector of polygenic score for a single genomic copy for all traits 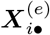 has a covariance matrix **Σ**^**(*X*)**^ under a Normal model:

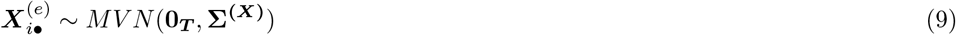

where *diag* 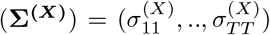 contains the variance explained by the polygenic scores of each trait, and the off-diagonal elements of **Σ**^**(*X*)**^ represent pleiotropic effects. For *C* full-genome copies we obtain a *C* × *T* matrix of polygenic scores with a matrix Normal distribution [19]:

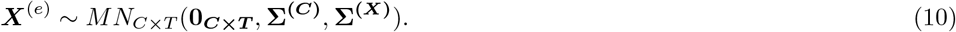

The matrix **Σ**^**(*C*)**^ represents (twice) the kinship coefficients between the *C* full-genome copies. For example, when the copies represent unrelated strains or lines, **Σ**^**(*C*)**^ = ***I***_***C***_; when they represent sibling embryos,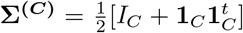. We assume that the chromosome scores are independent, with the scores matrix of each chromosome having the matrix Normal distribution:

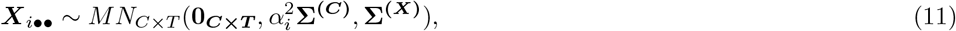

where 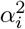 is the proportion of genetic variance explained by chromosome *i*. The genetic variances satisfy 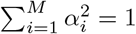, and this proportion is assumed to be the same for all traits, a consequence of the infinitesimal model and provided that the relative density of causal variants across the genome is similar across traits.

These contributions determine the utility of chromosomal selection, and are expected to be roughly proportional to chromosomes’ length or to their number of genes. Here, we show a numerical analysis with 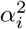 taken as the (normalized) chromosome lengths derived from the *S. cerevisiae* reference genome [8] and the human reference genome GRCh38 [37], shown in Table 2. The actual 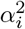 may vary across traits and can be estimated per trait by heritability-partitioning methods [15, 41] when needed.

Similarly, the non-score genetic components are modeled as:

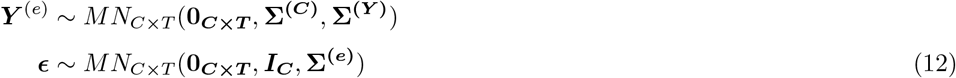

where diag **Σ**^**(*X*)**^ **+ Σ**^**(*Y*)**^ **+ Σ**^**(*e*)**^ = **1**_*T*_ , i.e. each trait is normalized to unit marginal variance while cross-trait covariances may be nonzero. The matrix **Σ**^**(*X*)**^ **+ Σ**^**(*Y*)**^ is known as the genetic covariance matrix, and can be estimated from GWAS data using e.g. methods like LD-Score-Regression [6]. The diagonal elements 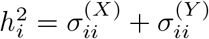 are the narrow-sense heritabilities of the traits.

The matrix **Σ**^**(*Y*)**^ **+ Σ**^**(*e*)**^ is the covariance matrix of the residuals, and determines the conditional distribution of the phenotypes vector conditioned on the scores vector. For simplicity, our model makes several standard assumptions: no shared environment (hence the identity ***I***_***C***_ is used as a covariance matrix for ***ϵ***), and no assortative mating. If these assumptions are violated, this can be encoded by the covariance matrices of our model.

#### Threshold traits

For threshold (e.g. disease) traits we use the liability-threshold model [14]. Trait *k* has prevalence *K*_*k*_ ∈ [0, 1] and liability 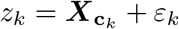, the selected score plus independent residual noise *ε*_*k*_ of variance 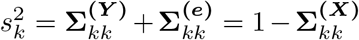 (the liability variance not explained by the score, with 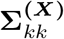 the score’s variance-explained on the liability scale).

The status is the binary random variable 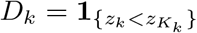 with threshold 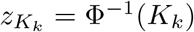 and **D** = (*D*_1_, … , *D*_*T*_). Conditioning on the score,

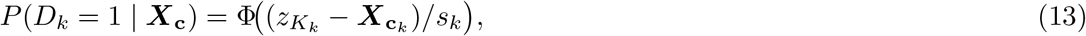

so a large score lowers the crossing probability; the all-pass probability *P* (**D** = **0**_*T*_ | ***X***_**c**_) is the Gaussian orthant probability of 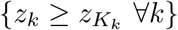 under the residual covariance **Σ**^**(*Y*)**^ **+ Σ**^**(*e*)**^ .

### C Implementation Details

Our algorithms and the simulation study presented in the paper are implemented in an R package available at https://github.com/orzuk/EmbryoSelectionCalculator, with the chromosomal-selection functions in code/R/chrom. Performance-critical routines are implemented in C++ via Rcpp [13].

## Footnotes

1 Partition: given positive integers *a*_1_, … , *a*_*n*_ with 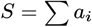, decide whether there exists *A* ⊆ [*n*] with 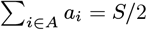. Partition is weakly NP-complete [17].

2 Our Branch-and-Bound implementation supports an optional heuristic cap of *B* retained Pareto-optimal vectors per stage (Appendix C); for all experiments reported in this section the cap was set to *B* = 10^7^ and was never reached, hence the reported solutions are genuine global optima of Problem 1.

